# TUSX: an accessible toolbox for transcranial ultrasound simulation

**DOI:** 10.1101/2021.06.03.446963

**Authors:** Ian S. Heimbuch, Guido C. Faas, Marco Iacoboni, Andrew C. Charles

## Abstract

Normally, the complicated nature of acoustic simulation makes it infeasible for most research groups doing individual transcranial ultrasound studies, hindering interpretation of results and complicating the determination of safety limits. We present here an open-source MATLAB toolbox to perform acoustic simulations using subject-specific medical images for transcranial ultrasound experiments. This toolbox, Transcranial Ultrasound Simulation Toolbox (TUSX), consists of an integrated processing pipeline that takes in structural MR or CT images, processes them for accurate simulation, and runs the simulations using k-Wave, an existing open-source acoustics package. We describe here the processing TUSX performs, along with its reasoning. We also validate its output using real-world pressure measurements in a water tank.

## Introduction

The use of ultrasound for non-invasive brain stimulation (NIBS) has been gaining increased attention in recent years, in part due to it offering increased spatial precision compared to other NIBS techniques. For example, a small single-element transducer produces an ellipsoid intracranial focus with a full width half maximum focus of 4 to 7 mm (Deffieux et al., 2013; Lee et al., 2015; Legon et al., 2014; Tufail et al., 2010). Since transcranial ultrasound can be so precise, accurate understanding of where its energy lands is crucial for interpretation of results. Reasonable success with tUS placement can be made with simple geometric targeting. However, skull bone significantly affects tUS pressure fields, sometimes leading to resulting pressure focuses deviating over a centimeter from the intended target (Hynynen & Jolesz, 1998; Lee et al., 2015).

Ultrasonic waves are acoustic waves above the frequency range of human hearing (>20 kHz). Ultrasonic waves mechanically propagate through a medium, with alternating periods of compression and rarefaction (stretching) as the pressure wave passes. The precise nature of this propagation depends on the characteristics of the medium, mainly the speed of sound, density, and attenuation coefficient (α), and the acoustic frequencies involved (Ono, 2020).

The mathematical nature of the interaction of particles interacting in an acoustic wave are well understood (Feynman et al., 1965). It is this understanding that allows for imaging uses of ultrasound such as sonography and non-destructive ultrasonic testing. As such, the large-scale nature of acoustic waves passing through materials can be modeled with various methods (Cox et al., 2007; Treeby & Cox, 2009). However, the options to perform such simulations are either commercial (e.g. multiphysics engineering simulation software) or accurate and open-source but with a learning curve. An example of the latter is k-Wave, a powerful open-source tool to perform computationally efficient acoustic simulations using a k-space corrected pseudospectral time domain (PSTD) scheme (Treeby & Cox, 2010). Since k-Wave is robust and flexible, it also leaves a novice user open to mistakes that can make implementation challenging or, worse, lead to erroneous results.

Here we present a toolbox that streamlines the process. Using this toolbox, TUSX (Transcranial Ultrasound Simulation Toolbox), simulations can be performed using conventional MRI or CT images with subject-specific precision. First, we outline the primary processing pipeline within the toolbox and discuss the reasoning for key features. Second, we show validation experiments that demonstrate the simulation results predict ultrasound pressure fields with sufficient accuracy. Finally, we discuss applications for this toolbox for transcranial ultrasound research in human participants.

## Methods

### Toolbox Overview

TUSX, which the authors pronounce as “tusks”, takes in a binary 3D skull volume (MR or CT). The skull volume is then scaled using interpolation to a higher resolution, which is necessary for accurate simulation. The location, trajectory, and other parameters of the ultrasound transducer is set by the user. The volume is then rotated such that the transducer trajectory is in line with the computational grid. This rotation improves accuracy by reducing the effect of ‘staircasing’ along the skull edge, which would otherwise cause aberrant interference patterns (J. L. B. Robertson et al., 2017). The skull volume is smoothed at multiple steps (before and after rotation) using morphological image processing (Figure 1).

**Figure 1.**
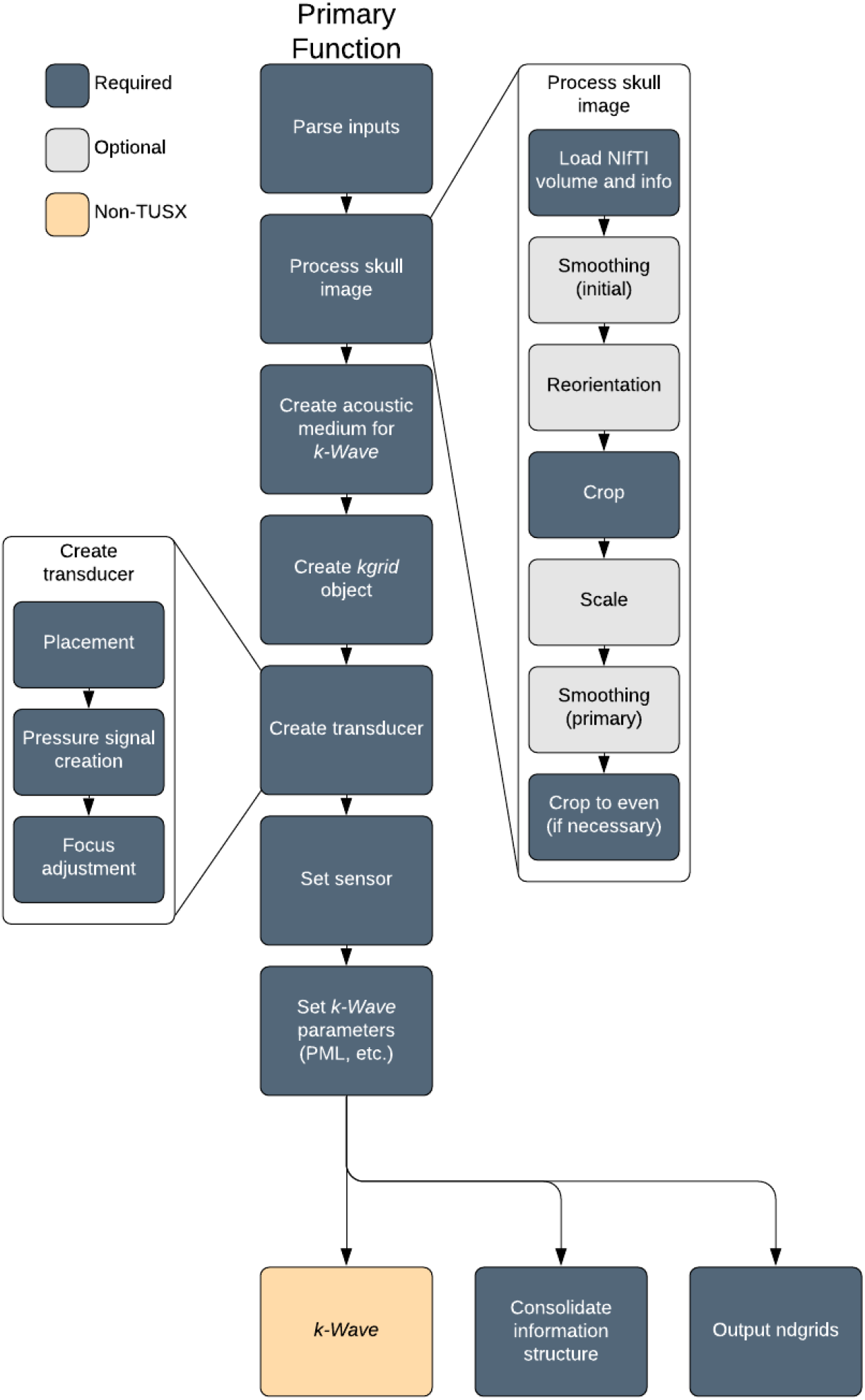
Diagram of primary TUSX pipeline. Primary Function) Major steps of the pipeline. Details for “Process skull image” and “Create transducer” steps are in breakout sections.

The proper acoustic parameters are then applied to each part of the volume. For MR images, a homogenous skull medium is used. For CT images, a homogenous skull medium or a heterogeneous skull medium derived from the apparent bone density is used (Connor et al., 2002). Additional k-Wave parameters are then set at proper values for the given volume. TUSX then executes the time domain 3D acoustic simulation via k-Wave (Treeby & Cox, 2010), which uses a k-space pseudospectral method. TUSX can execute the simulation via k-Wave’s MATLAB (Mathworks, Inc., Natick, MA) implementation or export it for use in k-Wave’s compiled C++ version for improved performance (Treeby et al., 2012).

#### Scaling

Due to the nature of the numerical methods used to perform the acoustic simulations, final accuracy depends on the spatial resolution of the simulation grid—specifically the ratio of simulation grid points per acoustic wavelength (PPW). Ideally, simulations would be performed at ~10 PPW or greater to assure no significant loss of accuracy (J. L. B. Robertson et al., 2017). But even for a high-resolution Human Connectome Project T1-weighted structural MRI (voxel width: 0.8 mm), simulating from the untouched volume gives a PPW of ~3.9 (using a tUS-typical 500 kHz source). However, by increasing the spatial resolution of the computational grid through interpolation of the volume, the PPW is raised to ~7.8 and ~15.6 for a 2X and 4X increase, respectively. TUSX uses nearest-neighbor interpolation when scaling skull masks (i.e. binary volumes) and linear interpolation when scaling CT bone density values.

Increasing the resolution used also has the benefit of allowing for more accurate representations of curved (or otherwise non-orthogonal) acoustic sources within the grid, which is also crucial for simulation accuracy (Wise et al., 2019).

#### Smoothing

The binary skull masks are smoothed by TUSX via morphological image processing in a multi-step process. First, morphological closing (i.e. dilation then erosion) is performed using a spherical structuring elements of radius 4 times the scaling factor (see Scaling). This fills in staircasing effects introduced by the nearest-neighbors interpolation of the initial binary skull mask. Second, morphological opening (i.e. erosion then dilation) is performed using a spherical structuring element of radius 1 times the scaling factor. This removes extraneous jagged protrusions from the skull mask, which are likely not present on the skull.

TUSX also has the option of performing an initial smoothing step before any other skull processing. This option involves morphological closing followed by opening with spherical structuring elements (radius: 1 grid point). While a relatively minor effect, this option is of particular use when reorientation is desired since the step smooths out minor protrusions and divots that could be accentuated by the linear transformation process.

As a whole, the smoothing process serves to assure an accurate representation of a human skull, since the curves of a typical cranium is fairly smooth. The process improves simulation accuracy since smoothing the scaled skull masks reduces the amount of staircasing along the curves of the skull, and such staircasing is a significant cause of simulation error (J. L. B. Robertson et al., 2017).

### Skull Mask Creation

#### Skull

TUSX creates a simulation volume by importing a medical image and applying the appropriate acoustic values to each part of the volume. NIfTI-formatted MR or CT volumes are supported for importation. Users can import a binary skull mask in which skull voxels have already been selected by a separate tissue segmentation program (e.g. BrainSuite or FSL BET).

For skull masks, the acoustic properties for bulk skull are applied to all points within the skull (i.e. no differentiation between compact and spongy bone). This approach allows for use of structural MRI for simulation, even though it does not have the same detail of bone density as CT. Importantly, the use of skull masks with homogenous skull acoustic properties has been shown to be effective in simulations within the frequencies used for transcranial ultrasound stimulation (Jones & Hynynen, 2016; Miller et al., 2015; J. Robertson et al., 2017). The acoustic properties to be used for bulk skull can be set by the user or kept to default TUSX values.

#### Brain

TUSX models the entire intracranial space as a homogenous medium, using the bulk acoustic properties of brain tissue. We chose this approach because it has been shown that modeling brain morphology does not significantly impact intracranial results (Mueller et al., 2016), due to the relatively minor differences in acoustic properties between intracranial media (i.e. grey matter, white matter, cerebrospinal fluid). As with skull, the user can set desired acoustic properties to be used for brain tissue or use default TUSX values.

#### Volume Dimensions

TUSX crops the simulation volume down to the region of interest to improve performance. Due to the spatially precise nature of tUS, many tUS experiments involve stimulation volumes significantly smaller than the full intracranial space. As such, TUSX can restrict the simulation to the area surrounding the tUS trajectory.

TUSX will also choose precise dimensions integers for best performance. Specifically, the computations performed by k-Wave involve fast Fourier transforms, which works best on powers of two or other integers with low prime factors (Treeby & Cox, 2010). TUSX executes this through a combination of removal of single slices to avoid odd integers, cropping, and the selection of the thickness for the perfectly matched layer (PML), which surrounds the volume to prevent waves from wrapping to the other side of the volume.

### Transducer Creation

Placement of the ultrasound transducer within the simulation grid is straightforward with TUSX. The user provides two coordinates in NIfTI voxel space: one for the center of the transducer face and one to set where the transducer is facing. The use of NIfTI voxel space significantly streamlines the process of placement for researchers, who are likely already working with NIfTI coordinates for other aspects of their projects.

An example workflow is as follows. 1) A researcher chooses a cortical ROI in their preferred imaging software (e.g. an fMRI hotspot or anatomical marker). 2) The researcher uses their preferred software to select a point on the scalp such that the transducer would face the ROI. 3) The scalp coordinate is placed into TUSX as the ‘transducer’ coordinate, and the ROI coordinate is input as the targeting coordinate.

Transducer size, shape, focal length, and acoustic frequency are also set by parameters provided by the user. TUSX creates the single-element transducer pressure source as a disc or curved spherical cap within the simulation grid (k-Wave function: *makeBowl*) (Ling et al., 2015). When a focused transducer is desired, TUSX performs beam forming by setting the pressure source to a spherical cap with a radius of curvature of the desired focal length. To improve focusing even further, the temporal delay of the sinusoidal pressure traces emitted from each source point are then adjusted to better match their aliased positioning on the orthogonal grid (k-Wave function: *focus*) (Martin et al., 2016).

### Validation: Water Tank

Pressure measurements were taken in an acrylic water tank filled with degassed deionized water. Measurements were made in a 3D volume surrounding the focus of the transducer pressure field. Measurements were done with two setups: with a skull analogue between the transducer and the hydrophone and with nothing between the transducer and the hydrophone (free water).

#### Ultrasound Equipment

A 500-kHz focused piezoelectric transducer (Blatek Industries, Inc., State College, PA) was used for all water tank measurements. The cylindrical transducer had a diameter of 3 cm and a focal point of 3 cm. The transducer was secured in a custom 3D-printed mount, which was affixed to the measurement rig. This transducer was chosen because it is representative of models used in various human transcranial ultrasound stimulation experiments (e.g. Legon et al., 2014; Heimbuch et al., 2021).

The transducer was driven by 500-kHz sine-wave voltage pulses from a waveform generator (33500B Series, Keysight Technologies, Santa Rosa, CA). Voltage pulses were amplified by a 50-dB radio frequency amplifier (Model 5048, Ophir RF, Los Angeles, CA). A 3-dB fixed attenuator was attached in line following the amplifier.

#### Measurement Equipment

Time-varying pressure traces were recorded via a hydrophone placed orthogonal to the face of the transducer (1 mm, Precision Acoustics Ltd, Dorchester, UK). The pre-amplified signals were sampled at 10 MHz by a PCI oscilloscope device (PCI-5105, National Instruments, Austin, TX), which were recorded using LabVIEW (National Instruments). Pressures were sampled across a three-dimensional volume using a micromanipulator, with multiple pressure traces sampled at each point on the sampling grid. Duplicate pressure traces were averaged offline in MATLAB to get a single average pressure trace per grid point per setup.

#### Skull Analogue

The skull analogue was a 4.6-mm thick polytetrafluoroethylene (PTFE) sheet (ASTM D3308) (Figure 2). We chose to use a flat analogue to create a best-case scenario for a one-to-one match between the real-world setup and its representation in simulation space—in this case an orthogonal grid. Specifically, the representation of a curve (e.g. a plastic analogue with the radius of curvature of a typical cranium) would introduce aliasing artifacts (see Smoothing).

**Figure 2.**
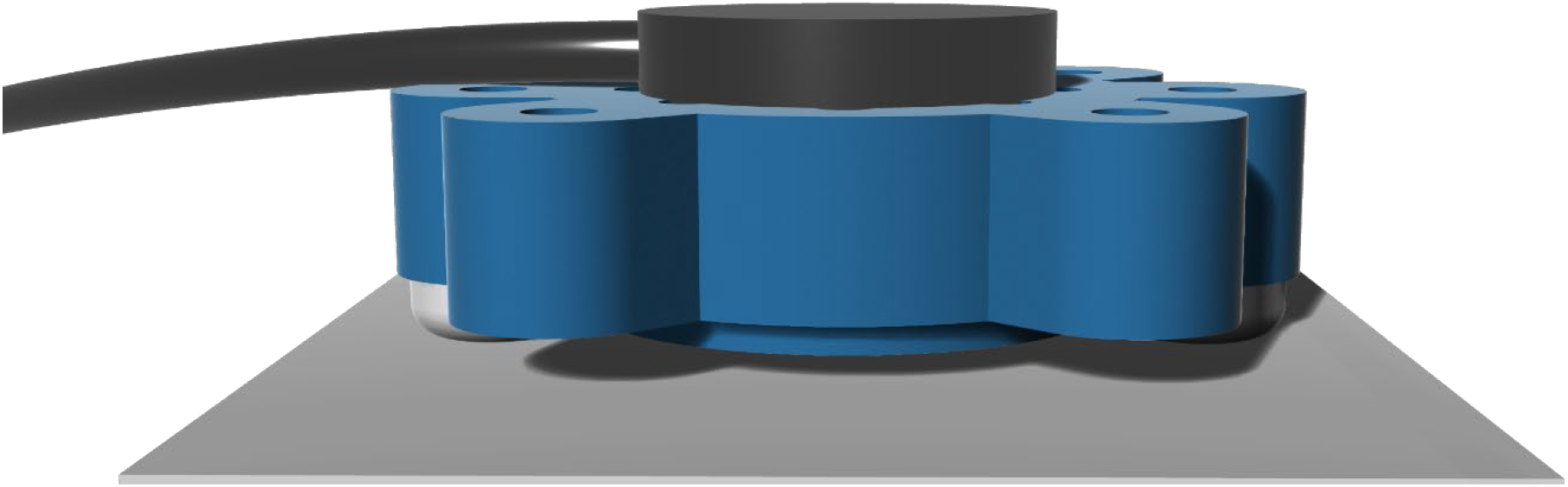
Render of transducer mount and skull analogue. The ultrasound transducer (black) sat in a 3D-printed housing (blue), which was mounted to a stereotaxic height control (not shown). For measurements with a skull analogue, the PTFE (white) was directly below the transducer mount, with a 4.8-mm gap between the transducer face and the PTFE. The PTFE was mounted to the same stereotaxic rig (not shown).

**Figure 0-4.**
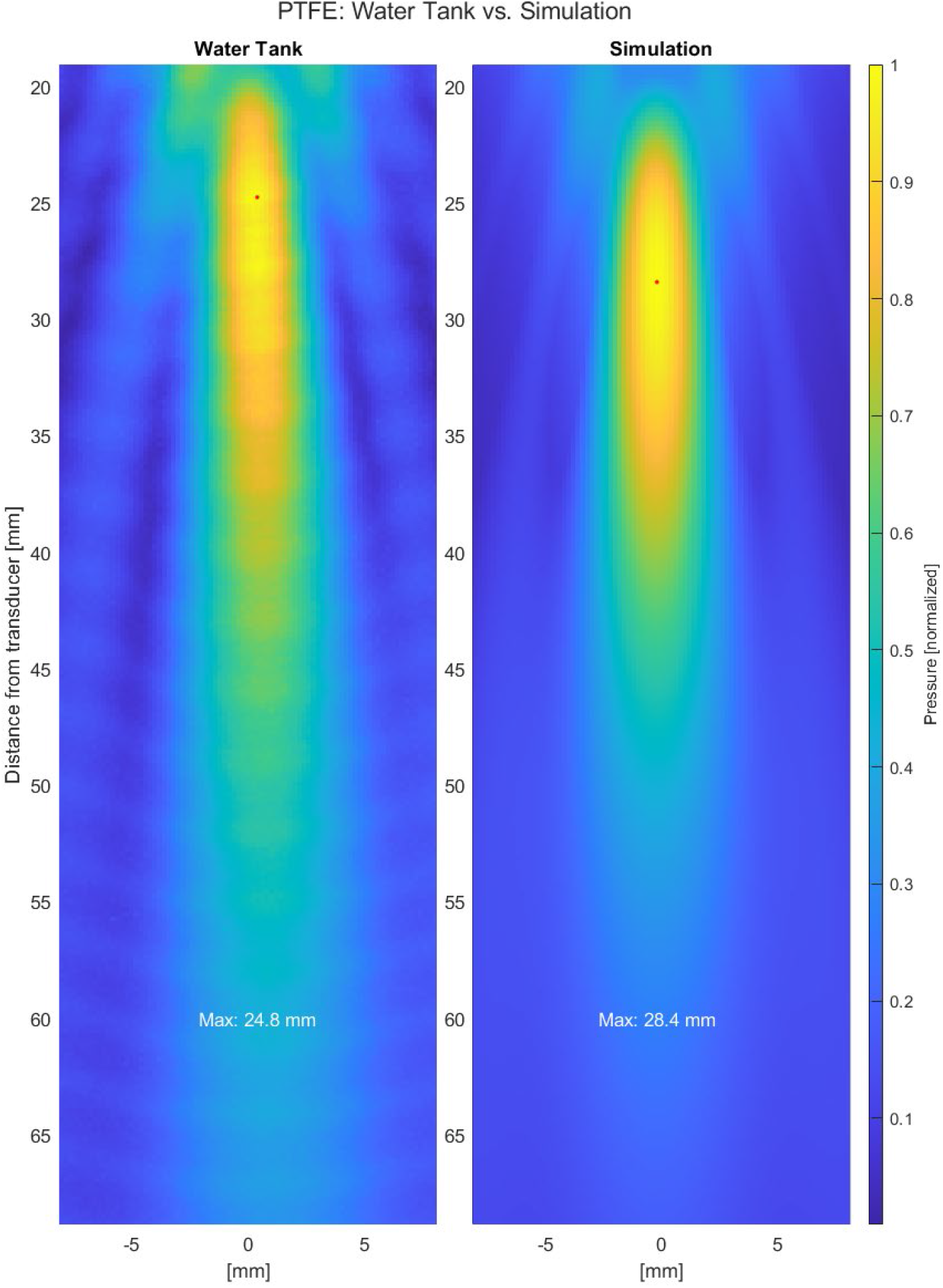
Real pressure distribution vs. simulated pressure distribution; PTFE. Left: A cross section of the pressure distribution of a focused ultrasound transducer in a water tank. PTFE skull analogue (4.6 mm thick) located below the tUS transducer. Right: A cross section of the pressure distribution of a focused ultrasound source simulation using the same medium properties of the water used in the water tank recording (Left) and the acoustic properties of PTFE. Simulated pressure source had a focal length of 32 mm. Both were measured and simulated, respectively, with grids point widths of 0.2 by 0.2 by 0.2 mm.

**Figure 5.**
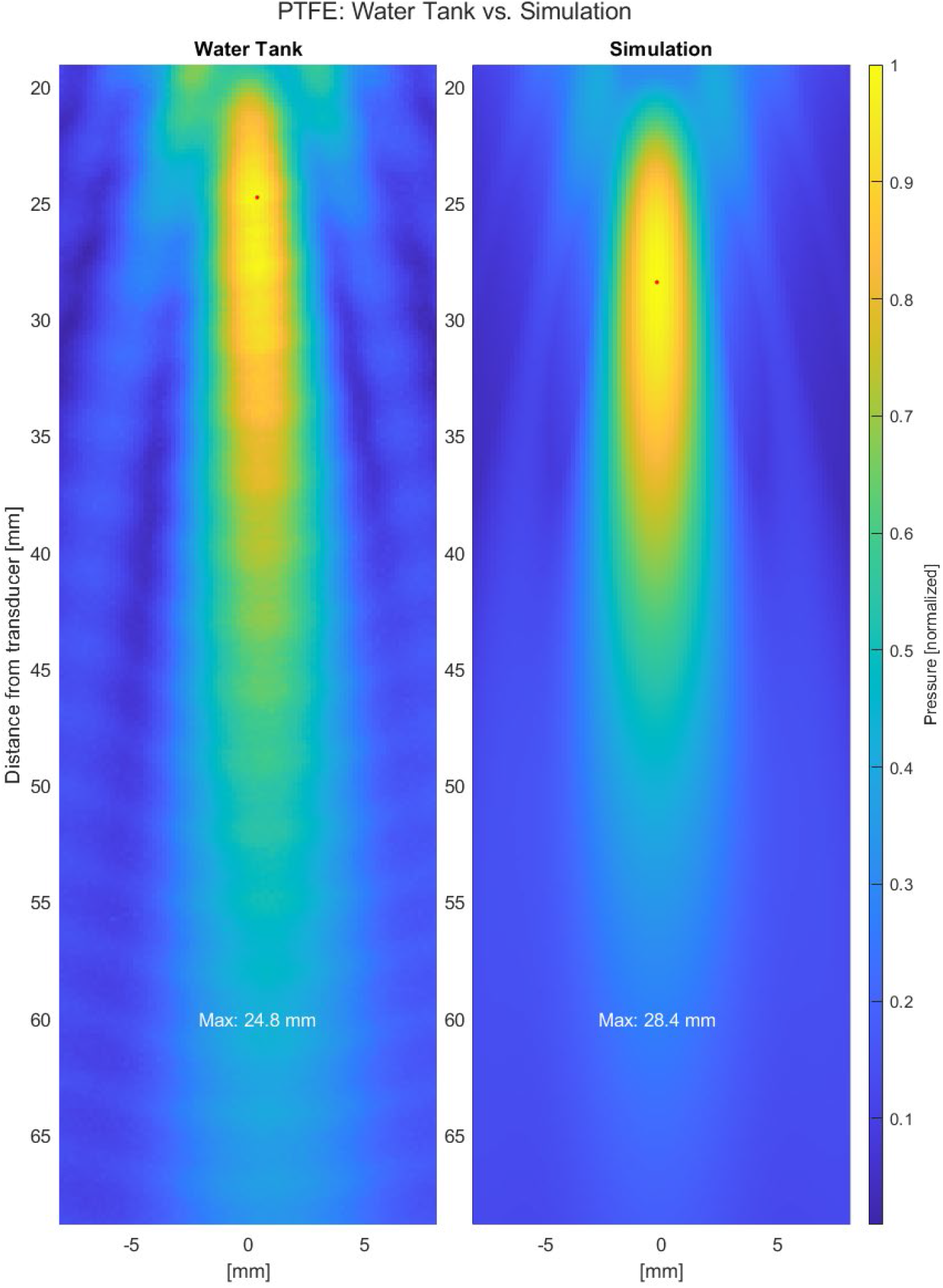
Real pressure distribution vs. simulated pressure distribution; PTFE. Left: A cross section of the pressure distribution of a focused ultrasound transducer in a water tank. PTFE skull analogue (4.6 mm thick) located below the tUS transducer. Right: A cross section of the pressure distribution of a focused ultrasound source simulation using the same medium properties of the water used in the water tank recording (Left) and the acoustic properties of PTFE. Simulated pressure source had a focal length of 32 mm. Both were measured and simulated, respectively, with grids point widths of 0.2 by 0.2 by 0.2 mm.

**Figure 6.**
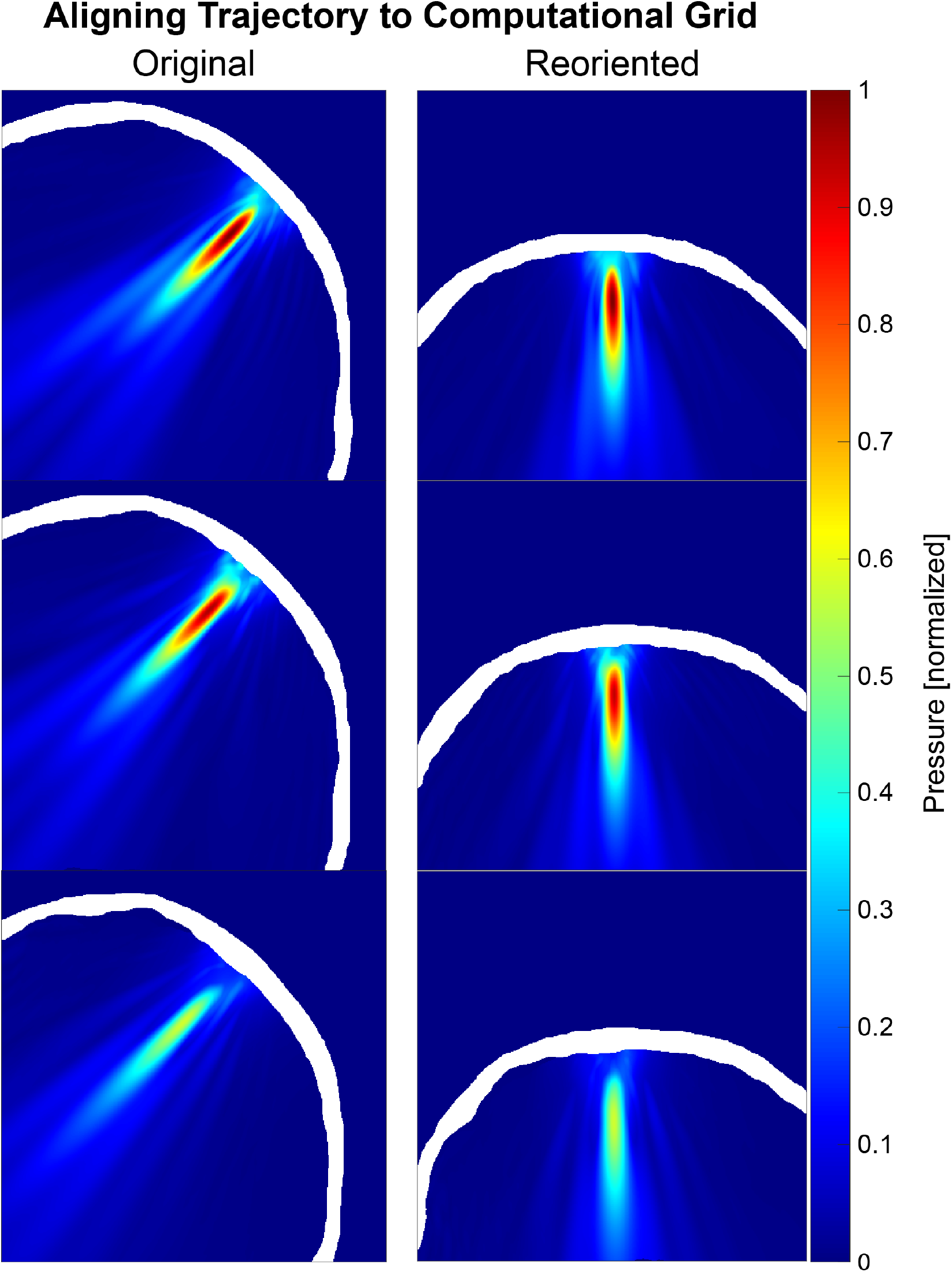
Simulation with and without alignment to computational grid. Three example parietal trajectories. Pressures are normalized to the maximum pressure for the respective simulation. In-brain pressures shown only. Left: Original volume. Right: Volume reoriented before simulation. Trajectory of the emitted ultrasound is now orthogonal to the 3D grid used for simulation.

To counteract curling caused by the fabrication process, the PTFE sheet was clamped between two steel plates and placed in a laboratory oven overnight. The PTFE was then supported with steel brackets. The PTFE was placed 4.8 mm from the transducer face.

#### Degassing System

The gas content of the water in the tank (~150L, ~55×55×50 cm) was kept low by continuously recirculating (~2 L/min) through a vacuum degassing system with a Liqui-Cell G453 2.5 × 8 Membrane Contactor (3M Separation and Purification Sciences Division, Charlotte, NC). The contactor was operated in the vacuum only mode. When water in the tank was replaced, the degassing ran at least for 20 complete exchanges (after ~25 hours) before beginning an experiment to ensure the water was completely degassed.

### Validation: Simulation

#### Simulation Grid

For validation experiments, acoustic simulations were performed via TUSX that matched conditions present in the real-world water tank measurements (Validation: Water Tank). Simulations were performed using a grid spacing of 0.2 by 0.2 by 0.2 mm, which matched the highest spatial resolution used for water tank measurements. A NIfTI mask volume (dimensions: 236 × 236 × 492 voxels) that matched the setup of each tank experiment was fed into TUSX in place of an MR or CT head volume (i.e. a 4.6-mm thick mask layer for the PTFE experiment and an empty volume for the water experiment). PML was set to 10 grid points, creating a final simulation grid of 256 by 256 by 512 grid points.

The pressure source for the validation experiment was set to mirror the transducer used in the water tank. This consisted of a focused source with a 32-mm focal length at 500 kHz (see Transducer Creation), with the face of the transducer oriented orthogonal with the computational grid. A focal length of 32 mm, rather than the manufacturer specification for the transducer of 30 mm, was chosen to better match the water tank data when measured in free water (Supplemental Figure 2). Simulations were performed at a temporal interval of 285 temporal points per period (PPP) for a Courant-Friedreichs-Lewy (CFL) number of 0.0519, which is well within guidelines for accurate simulation using k-Wave’s pseudospectral time domain scheme (J. L. B. Robertson et al., 2017).

#### Acoustic Properties for Validation

Speed of sound of the water in the tank at the time of each recording was calculated by the mean delay arrival time between sequential sample points in the Z axis (away from the transducer) (~1502 m/s), since the distance between sample points was known (0.4 mm for speed of sound measurements). These calculations were with mean pressure traces taken from multiple sequential samples, and delays were found using cross-correlation with the input voltage trace. Density and attenuation were set to referenced values (density: 998 kg/m^3^; 2.50×10^−5^ dB/cm) (Duck, 1990).

Speed of sound of the PTFE was set using a referenced value: 1310 m/s (Ono, 2020). Density and attenuation of PTFE were set to manufacturer specifications and a referenced value, respectively (density: 2131 kg/m^3^; 7.36 dB/cm) (Ono, 2020).

### Validation: Comparison

#### Alignment

Comparison between volumes of measured water tank pressures and simulated pressures were enabled by aligning the two grids based on distance from the ultrasound transducer (real or simulated). Spatial calibration for the water tank measurement grid was performed by calculating the distance of central grid points from the delay between signal and pressure onset, since the speed of sound was known (see Acoustic Properties for Validation).

#### Error Metrics

In addition to visual inspection of aligned slices, comparisons between water tank and simulation results were aided with two error metrics: spatial deviation of the focus and the full width at half maximum (FWHM) of the focus. Specifically, we used the center of mass of the −3 dB focus area (i.e. center of mass above ~79% of the maximum of the 3D volume) and the FWHM in the X- and Y-planes averaged.

## Results

### Water Tank Validation

#### Free Water

We compared the results of a 3D acoustic simulation prepped through TUSX that matched the setup of a 3D pressure measurement tank in free water. Results show that the simulation produced a pressure field that closely matches the real-world pressure field (Figure 3). The center of mass of the −3 dB focus of the real-world transducer was at a depth of 29.6 mm, while the parameter-matched simulation showed a center of mass at 30.1 mm (delta: 0.48 mm). The FWHM of the real-world transducer focus was 4.6 mm, while the simulated focus had a FWHM of 4.0 mm.

#### Skull Analogue

We also compared a parameter-matched simulation to a water tank measurement in which a flat skull analogue (PTFE, 4.8 mm) was placed against the transducer. This comparison also showed a close match between simulated and measured pressure fields. The center of mass of the −3 dB focus of the ‘transcranial’ pressure field was at a depth of 30.5 mm, while the parameter-matched simulation showed a center of mass at 30.8 mm (delta: 0.29 mm). FWHMs mirrored those in free water (water tank: 4.6 mm; simulation: 4.0 mm). In the water tank pressure field, we did observe some deviation of the pressure field from the Z axis. This is likely due to residual warping in the PTFE and/or imperfect alignment of the transducer by the 3D-printed mount.

## Discussion

We have created a MATLAB toolbox, TUSX, that streamlines the process of performing acoustic ultrasound simulations with subject-specific medical images for transcranial ultrasound experiments. In addition to detailing the processing steps performed by TUSX to enable accurate simulations, we have validated the accuracy of TUSX (and the acoustic simulation package it uses, k-Wave) via real-world measurements in a water tank.

### Advantages of TUSX

#### CT vs. MRI for Transcranial Ultrasound Simulation

The support for use of MRI for acoustic simulations, over simply CT, significantly broadens the accessibility of TUSX. CT is the standard for calculation of phase correction in high-intensity focused ultrasound (HIFU) for tissue ablation, but the desire to avoid radiation exposure for healthy volunteers makes the acquisition of MRI a more appealing option—though CT has been used for acoustic simulation in two non-clinical tUS studies (Lee et al., 2015, 2016). Beyond radiation considerations, MRI is more feasible to acquire, since many neuroscience labs have existing MRI access and experience. While MR cannot directly derive bone density as CT can, MRI has already been validated as an adequate reference for transcranial focusing of ultrasound (Hynynen & Sun, 1999; Miller et al., 2015; J. Robertson et al., 2017; Wintermark et al., 2014), with results approaching that of CT. Especially considering that the margin of error for tUS is likely higher than HIFU aiming to ablate precise tissue volumes (e.g. specific thalamic nuclei), we believe the use of MRI provides adequate accuracy for acoustic simulations for tUS studies.

#### Reorientation

The feature to realign the skull volume in TUSX such that the ultrasound trajectory is orthogonal to the computational grid should provide for improved, consistent results for tUS targets that necessitate trajectories at oblique angles. This is because reorientation of the volume, along with subsequent smoothing, avoids having the ultrasound pressures interface with the skull at aliased portions of its curvature, and such ‘staircasing’ has been noted as the most serious cause of error in previous investigations of acoustic simulation accuracy (J. L. B. Robertson et al., 2017). As an additional tangential benefit, reorientation to an orthogonal trajectory could also improve the performance of the PML, since the PML performs better on waves of low angle of incidence (J. L. B. Robertson et al., 2017).

#### 3D Simulations

We chose to support 3D simulations with TUSX, compared to 2D simulations using slices through a 3D volume, for two reasons. First, simulating in 3D should provide a more accurate representation of the real pressures and exposures created during tUS. Second, the preparation of accurate 3D simulations has an additional degree of complexity compared to 2D. As such, a tool for 3D simulations should hopefully provide the most help to researchers.

#### Ease of Use

For a novel user performing simulations of transcranial ultrasound, TUSX greatly simplifies performing k-Wave acoustic simulations compared to k-Wave alone. It saves the user significant labor processing image volumes, placing ultrasound sources, and researching the necessary acoustic and k-Wave parameters for accurate transcranial simulations.

## Conclusions

Since skull morphology varies highly between individuals, variations in skull thickness and shape result in equally varied intensity levels and foci location following skull transmission of ultrasound. As such, having access to estimations of in-tissue intensities in each research participant is helpful for both safety and interpretation of results. TUSX lowers the high barrier to entry for transcranial ultrasound researchers to include acoustic simulation in their projects.

tUS is still a novel field, and the field is still developing a collective understanding of what neural responses can consistently be elicited by tUS and what tUS parameters it takes to do so. This need to synthesize optimized tUS parameters has been increasingly brought forward by members of the field in the past few years. Our hope is that by enabling more frequent reporting of in-brain tUS pressures and locations, we can more rapidly move toward that goal.

## Supplementary Figures

**Supplemental Figure 1.**
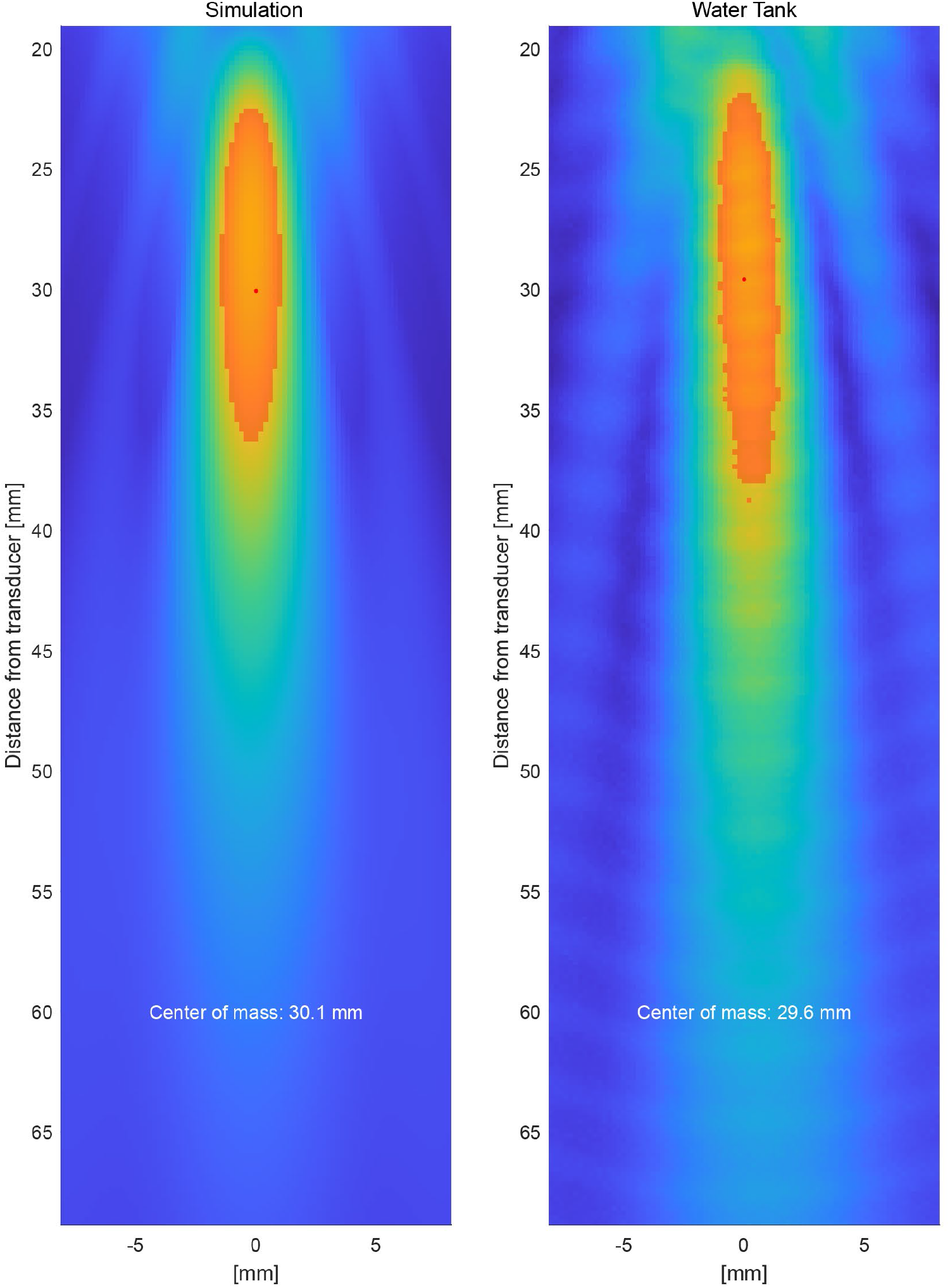
Real pressure distribution vs. simulated pressure distribution in free water; with center of mass. Orange overlay: −3 dB focus area (i.e. volume above ~79% of the maximum of the 3D volume). Red dot: center of mass of the −3 dB focus area in 3D space. Left: A cross section of the pressure distribution of a focused ultrasound transducer in a water tank. Right: A cross section of the pressure distribution of a focused ultrasound source simulation using the same medium properties of the water used in the water tank recording (Left). Simulated pressure source had a focal length of 32 mm. Both were measured and simulated, respectively, with grids point widths of 0.2 by 0.2 by 0.2 mm.

**Supplemental Figure 2.**
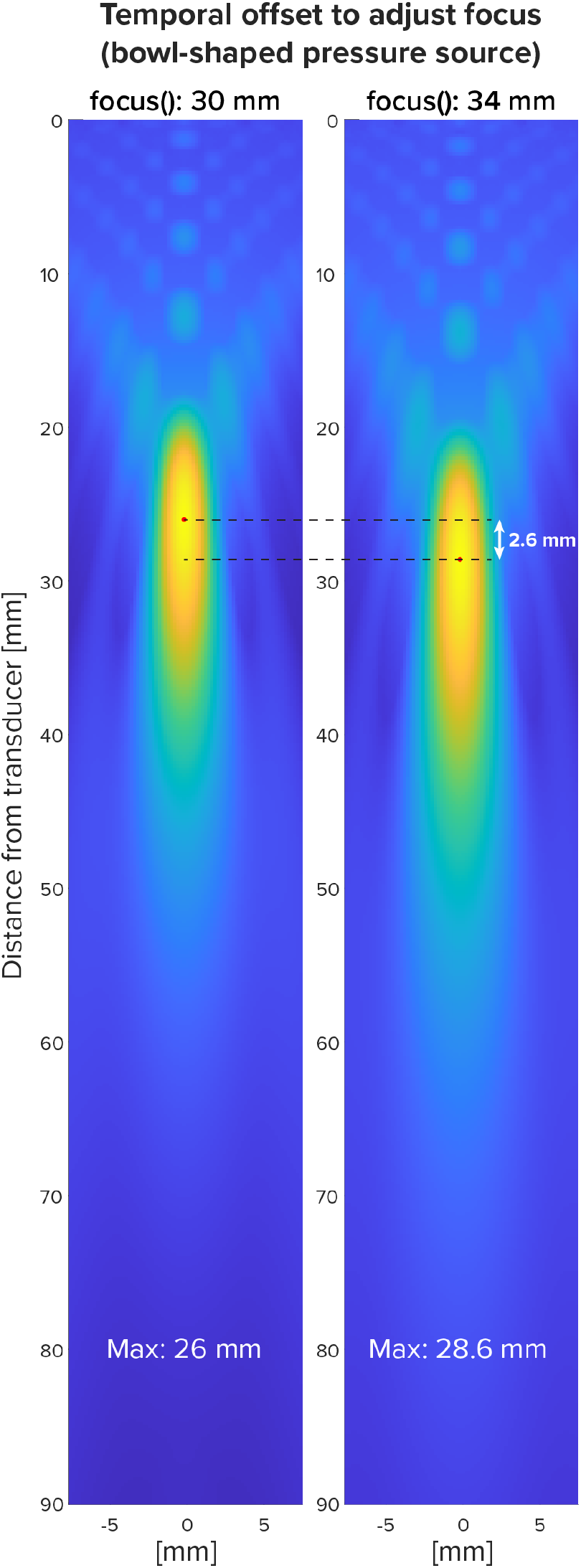
Adjusting focus with point-specific pressure trace delay. Comparison of two simulations in which the pressure source is a spherical cap with a 30-mm radius of curvature. Left: Same sine wave emitted from all points of spherical cap source (i.e. a pressure focus set to match the 30-mm radius of curvature). Right: Sine waves onset is based on the euclidean distance to focal point, now set to 34 mm. Distances [mm] are relative to the furthest point of the spherical cap (i.e. back). Temporal adjustment was made using the k-Wave function ‘focus’. Acoustic values of medium for simulation were that of brain.

## Acknowledgements

The authors would like to thank Mayank Jog for insightful comments on the manuscript.

## Notes

https://www.tusx.org

### Competing Interest Statement

The authors have declared no competing interest.

